# The relationship between microbial community succession, decay, and anatomical character loss in non-biomineralised animals

**DOI:** 10.1101/2024.07.01.601543

**Authors:** Thomas Clements, Robert Goodall, Sarah Gabbott, Duncan Murdock, Martha Clokie, Andrew Millard, Christopher Turkington, Orla Bath-Enright, Mark Purnell

**Affiliations:** GeoZentrum Nordbayern, Department of Geography and Geosciences, Friedrich-Alexander Universität Erlangen-Nürnberg, Erlangen, Germany; University of Leicester, Centre for Palaeobiology & Biosphere Evolution, School of Geography, Geology and the Environment, University Road, Leicester, LE1 7RH UK; Oxford University Museum of Natural History, Parks Road, Oxford OX1 3PW, UK; University of Leicester, Department of Genetics and Genome Biology, Leicester, UK; APC Microbiome Ireland and School of Microbiology, University College Cork, Cork, Ireland; Staatliches Museum für Naturkunde Stuttgart, Rosenstein 1, D-70191, Germany; School of the Environment, Geography, and Geosciences, University of Portsmouth, Portsmouth, Hampshire, PO1 3QL, UK

**Keywords:** Taphonomy, soft tissue preservation, decay, microbiome, fossilisation, fossil, bacteria

## Abstract

A fundamental assumption of hypothesis-driven decay experiments is that, during decay, the loss of anatomy follows a sequence broadly controlled by the intrinsic compositional properties of tissues. Recent work investigating the succession of postmortem endogenous microbial communities (thanatomicrobiome) challenges this assumption. These studies suggest the thanatomicrobiome exhibits a predictable, clock-like succession in response to physical and chemical environmental changes within a carcass. Therefore, it is possible that reproducible sequences of character loss during decay are controlled by thanatomicrobiome succession dynamics. If so, exceptionally preserved fossil anatomy would reflect a succession of ancient contemporaneous microbial communities, about which we know nothing, rendering decay experiments uninformative. Here, we investigate two questions: (1) what is the role of exogenous and endogenous bacteria during formation of the thanatomicrobiome and (2) do thanatomicrobiome successions control the sequence of anatomical character loss within a decaying carcass? Our analysis shows that the thanatomicrobiome is dominated by endogenous bacteria and that, even in the presence of inoculum, exogenous bacteria do not invade the carcass and replace native bacteria (while the carcass is intact). This confirms that the use of environmental inoculum in decay experiments introduces an inadvisable confounding variable. Secondly, we find no correlation between thanatomicrobiome successions and the sequence of anatomical character loss, supporting that fossil non-biomineralised characters correlate with their propensity to decay in extant relatives. These findings indicate that the inability to model ancient bacteria does not invalidate decay experiments. We also present a synthesis of the role of bacteria in non-biomineralised fossilisation.

## Introduction

Understanding the taphonomic processes that affect organic remains postmortem is vital when interpreting fossil organisms, especially those that preserve highly informative non-biomineralised anatomy. Decay is perhaps the dominant control on fossilisation of organic remains; the combination of autolysis and microbial metabolism acts from the moment of death and, under normal conditions, will proceed until all organic remnants are obliterated, preventing fossilisation from occurring. In amenable environmental conditions, processes that promote preservation – maturation and mineralisation – act to disrupt decay and stabilise organics over geological timescales to create fossils. Thus, to accurately interpret enigmatic non-biomineralised organisms from deep time, it is essential to understand that decaying organic remains are the substrate that preservational processes act upon during fossilisation, and that these remains will never be completely intact. Consequently, investigating the process(es) of decay is of primary concern of palaeontologists (e.g. Briggs and Kear, 1993a, b; Sagemann et al. 1999; Donoghue and Purnell, 2009; Purnell *et al*. 2018; Gabbott *et al*. 2021; Clements *et al*. 2022).

Decay experiments – hypothesis-driven experimental taphonomic investigations – are one of the most invaluable tools for analysing of the decay of non-biomineralised tissues and gross-anatomy. Robust taphonomic investigations are typically lab-based and are designed to investigate specific aspects of decay, biostratinomic processes, environmental variables that impact decay, or individual preservation processes. It should be noted that laboratory simulations that aim to replicate ancient environments or recreate the outcomes of fossilisation fall outside the scope of experimental taphonomy, as the number of unconstrained and confounding variables typically involved in such simulations renders their results open to multiple equally plausible interpretations (equifinality; see Purnell et al. 2018 for a review of experimental design). In the last few decades, taphonomic experiments have significantly furthered our understanding of the preservation of non-biomineralised remains. With each experimental result, our understanding of key concepts improves: for example, transport and rapid burial mechanisms (e.g. Bath Enright et al. 2017), environmental controls on rate of decay (e.g. Briggs and Kear, 1993a), how microbial decay drives the formation of geochemical conditions favourable for mineralisation (e.g. Sagemann *et al*. 1999; Clements *et al*. 2022), and that sediment composition plays an important role in maintaining the integrity of carcasses (e.g. Wilson and Butterfield, 2014). However, one of the most important observations revealed by experimental work is that during decay non-biomineralised anatomical characters are lost from the carcasses of organisms in an irreversible and reproducible (non-random) sequence (e.g. Sansom et al 2010, 2011; Murdock et al 2014; Nanglu et al 2015; Redelstorff and Orr 2015; Sansom 2016; Beli et al 2017). This is a fundamental concept as the procession of decay-modified characters are the anatomical substrate upon which stabilisation and preservation processes can act, and the non-random nature of decay allows taphonomists to model and predicted the continuum of character loss. Combining these models with our current understanding of the timing, interplay, and selectivity of information stabilisation (preservational processes) across soft tissues, we can begin to model preservational regimes (see Gabbott *et al*. 2021). Furthermore, as sequences of anatomical character loss have been demonstrated to be conserved between related organisms, combining the results of decay experiments with our understanding of preservational processes can shed light on the decay of fossil taxa which share homologous anatomical characters, a vital tool for interpreting enigmatic organisms (e.g. Sansom *et al*. 2011; 2013). Currently, the experimental approach is the only method that can generate detailed anatomical character decay sequence data.

Character-focussed taphonomic experiments from which decay sequences can be derived are relatively recent. Historically, observing and describing morphological decay has been a staple of decay experiments (e.g. Allison 1988a; Briggs and Kear 1993, 1994 a, b), the newer character-based methodology is increasingly useful because it provides data that is amenable to statistical analysis and allows investigation of systematic biases (e.g. Sansom et al 2010, 2011; Murdock et al. 2015; Cavicchini et al. *in review*). Importantly, character-focussed decay experiments have confirmed the underlying reproducible pattern of information loss outlined in older taphonomic work, generally in terms of ‘stages of decay’. In many previous studies, it was generally assumed that soft tissues were lost to decay in a sequence determined by the relative resistance of different biomolecules to autolysis and decay (Allison, 1988b; Tegelaar et al 1989; Gill-King et al. 1997; Briggs, 1999; Briggs et al. 2000; Briggs 2003). Briggs (2003), for example, listed the sequence of organic materials as decreasing in their degrees of resistance to decay as follows: macromolecules > lipids > carbohydrates > proteins > nucleic acids (with some caveats; based on Tegelaar *et al*. 1989). At a gross level, this assumes that the intrinsic biochemical properties of soft tissues control their resistance to decay. However, experimental work has shown that reality is more complex: it is not possible to accurately predict sequences of character loss from composition alone. For example, lampreys possess several different anatomical characters composed of cartilage, but these differ in their resistance to decay and are not all lost at the same point in the decay sequence (Sansom et al. 2011, 2013). Notwithstanding the important subtleties of the sequence of character loss revealed by decay experiments, few taphonomists would argue against the view that the biochemical properties of tissues provide the fundamental control on their propensity to decay.

However, recent work within the field of microbial forensic science presents a potential challenge to this hypothesis and an alternative explanation for the repeated sequences seen in anatomical decay. Studies on humans, pigs, and mice, in laboratory, terrestrial, and aquatic environments suggests that a distinct internal postmortem microbiome forms postmortem, termed the thanatomicrobiome (i.e., *thanatos*-, from Greek; death microbiome; Can et al. 2014), which colonises and spreads through a carcass in a predictable, clock-like succession based on changes in the physical and chemical environment (Dickson et al. 2011; Pechal et al. 2013; Metcalf et al. 2013, 2016; Can et al. 2014; Cobaugh et al. 2015; Guo et al. 2016; Lauber et al. 2016; Javan et al. 2016, 2019; Deel et al. 2020; Zhou et al. 2021; Aragonés et al. 2022). The formation of the thanatomicrobiome is dynamically influenced by both the *ante mortem* internal (endogenous) microbiome and by exogenous bacteria that live either on the carcass or in the surrounding environment (e.g. Dash and Das, 2020). This raises the possibility that the reproducible patterns and sequences of anatomical decay seen in taphonomic experiments are controlled by microbial succession, potentially by various bacteria taxa specialising on different biomolecules and tissue types. This hypothesis, unless it can be rejected, poses a significant challenge to experimental analysis of the role of decay in fossilisation, as rather than reflecting the nature of exceptionally-well preserved fossil organisms themselves, soft tissue fossils would actually represent the succession of contemporaneous ancient microbial communities, about which we know nothing (and are unlikely to ever be able to model).

Here, we address two fundamental questions. First, we address the question of the importance of exogenous and endogenous bacteria in the formation of the thanatomicrobiome within a carcass. It is possible that non-native exogenous microbial species invade decaying carcasses, and if they do so, they may dominate and/or replace the endogenous microbial communities. This question has a bearing on both the potential controls on decay – how important is the microbiota of the environment in which an organism decays – and the design of decay experiments. Although not previously addressed in the context of experimental taphonomy, the idea that exogenous bacteria are important provides the implicit methodological justification for the use of microbial inocula in many previous analyses of decay (e.g. Briggs and Kear, 1993a; Briggs et al. 1993; Hof and Briggs, 1997; Sagemann et al. 1999; Duncan et al. 2003; Martin et al. 2004; McCoy et al. 2015). Despite the widespread use of inocula, investigators generally know little about the community structures of the introduced microbiota or its role in subsequent decay.

Second, we address the key question outlined above: the hypothesis of correlation between the succession of the bacterial communities that develop through time as a carcass is broken down, and the sequence of loss of anatomical information.

## Methods

The questions outlined above, expressed as null hypotheses, provide the structure for our experimental methods. This study was run in conjunction with an experiment to test hypotheses regarding the relationship between substrate entombment and character loss (Murdock *et al*. in prep) and thus our methods are shared with this study.

We selected amphioxus (*Branchiostoma lanceolatum*, phylum Chordata) as the organisms for this investigation. As close living relatives of early vertebrates, amphioxus have been studied in several experimental decay investigations (Briggs and Kear, 1994b; Sansom et al., 2010; 2011; 2013, Murdock et al. *in prep*). The amphioxus were euthanised by overdose of tricaine methanesulphonate (MS222; 2 mg ml^-1^ with buffer), which has been shown to not adversely affect bacterial populations (see Sansom et al. 2010 and refs therein) in compliance with UK Government Guidance on the Operation of Animals (Scientific Procedures) Act, 1986, v.2014.

The null hypothesis regarding exogenous and endogenous bacteria is that the communities that develop within a decaying carcass through time are dominated by endogenous taxa, i.e. they come from within the carcass itself (hypothesis 1). For this experiment, individual euthanised amphioxus individuals (*n* = 10) were placed in clear polystyrene boxes (48 cm^3^) containing artificial seawater (ASW; Tropic Marin mixed with distilled water; salt content 33–36 ppm, pH 8). An inoculum was then created using a sediment slurry collected from an estuarine setting. In many previous decay experiments, bacterial inoculums taken to be ‘representative’ of natural environments were also cultured from estuarine sediments; this is because of the elevated rates of organic matter degradation by both aerobic and anaerobic bacterial respiration and the reported salinity tolerance of estuarine bacteria were thought to confer properties that are also useful to degradation (e.g. Briggs and Kear, 1993a). In these previous decay experiments, sediment and water mixtures have been collected from the Tay estuary, Scotland (e.g. Briggs et al. 1993; Briggs and Kear, 1993a; Hof and Briggs, 1997; Briggs and Kear, 1994a, b) and the Tamar River estuary, England (e.g. Sagemann et al. 1999; Duncan et al. 2003; Martin et al. 2004). We collected from the Tamar River estuary, at Weir Quay, Devon. A small trench was dug close to the low tide mark, and seawater and sediment (that presented as anoxic based on the clear transition from brown to black sediment colour, accompanied by a strong sulphur smell) was collected. This sediment sludge was mixed at 50 ml per litre ASW as per Briggs and Kear (1993a) and Martin et al. (2005). The polystyrene containers were sealed using silicon grease (Ambersil M494) to prevent gas exchange and invasion of bacteria external to the experiment. The containers were incubated at 25°C in a maturation chamber for the duration of the experiments and were removed at approximately logarithmically spaced intervals (days 1, 28, 134, and 1138). For each sampling point there was one control (decayed in ASW without inoculum) and three inoculated amphioxus. At each sampling point, morphological character presence/absence data were collected from each amphioxus carcass using microscopy, and then each sample was destructively sampled for DNA extraction. Amphioxus organic material was removed from the polystyrene container, placed within a 2 ml microcentrifuge tube, and suspended in an amount of lysis buffer (Tris-1mM EDTA buffer [pH 8.0] containing 5mg/ml proteinase K and 10% (w/v) Sodium dodecyl sulfate) needed to create a 2 ml sample. The sample was then vortexed until fully homogenised and flash frozen at - 80°C and stored until required.

The null hypothesis regarding bacterial community succession and anatomical decay is that the communities that develop within a decaying carcass through time are not correlated with the sequence of character loss (hypothesis 2). The experiment was designed to test this hypothesis alongside an analysis of the relationship between character loss and sediment entombment (Murdock et al. in prep), with samples of amphioxus decayed in three different sterile substrates: Kaolinite (Puraflo China Clay, Clayman Supplies Ltd.), Mica (muscovite, 2 µm to 1 mm, Imerys Ceramics), and Bentonite (predominantly montmorillonite from Wyoming, Clayman Supplies Ltd.). The substrates were mixed with the ASW (Tropic Marin mixed with distilled water; salt content 33–36 ppm, pH 8) to form a sludge, each approximately equal in consistency and density (clay by weight: ASW - kaolinite, 3:4 = 1.5 g/cm^3^; mica, 6:5 = 1.6 g/cm^3^; bentonite, 2:3 = 1.4 g/cm^3^). The slurries were then poured into clear polystyrene boxes (48 cm^3^), filling them to the halfway mark. A stainless-steel mesh was laid on top of the sludge (to aid carcass exhumation at the sampling interval) and a freshly euthanised amphioxus was placed upon the mesh. The container was then filled with the respective sludge, entombing the carcass completely, and then incubated at 25°C for the duration of the experiment. Three samples of each substrate type were removed from the incubator at the following sampling points: days 2, 8, 21, 28, 35, 42, and 134.

Exhumation of samples entombed within a substrate presents a challenge, as removal can cause damage to the carcass. Each sample was flash frozen in liquid nitrogen immediately after the containers were removed from the incubator. The then frozen substrate block (with entombed carcass) was carefully removed from the polystyrene box. In sterile conditions, the substrate block was cracked open along the plane of the stainless-steel mesh, exposing the overlying carcass, which was then examined to identify morphological character states for Murdock *et al*. (in prep). The underlying substrate block was kept frozen. From this, approximately 1 mm of substrate closest to the carcass along with any persisting organic material was sampled and placed in a sterilised beaker of lysis buffer, producing a slurry. As the experimental setup was sterile, any bacteria found in the slurry must be derived from the carcass. This slurry was vortexed and poured into a 2 ml microcentrifuge tube for DNA extraction and then flash frozen and stored until analysis. DNA extraction was performed on 53 viable samples across all substrates (see supplemental table 1.1). DNA extraction methods are outlined below. It should be noted that there was a high failure rate in DNA extraction, resulting in 10 successful extractions (see experimental limitations below).

### Bacterial DNA extraction, amplification, and sequencing

All DNA extraction, purification, and sequencing was performed in the Department of Infection, Inflammation, and Immunity laboratories at the University of Leicester. The samples were thawed and DNA extraction was carried out using the method appropriate for the sample type (e.g. sediment, tissue, solution) using either phenol–chloroform–isoamyl alcohol method as outlined in Nale *et al*. (2015), Qigen DNeasy Blood and Tissue kits, and DNeasy PowerMax Soil kits (see supplemental table 1.1). At the qPCR stage, kit controls and negative controls were run to test for any potential contamination in the methodology; any values returned were subtracted from the final qPCR results accordingly. Once DNA was extracted from the samples, the 16S rRNA gene region was sequenced using Illumina HiSeq sequencing, a standard practice for taxonomic characterisation of bacteria (e.g. Imam *et al*. 2019). The sequencing protocols used in this study followed the Illumina protocol for 16S Metagenomic Sequencing Library Preparation Revision B (Amplicon *et al*. 2013). Primers used targeted V3 – V4 region of the 16S ribosomal RNA gene and can be found in supplemental table 1.2. DNA sequencing allows bacteria within the samples to be categorised as operational taxonomic units (OTUs), and we utilised the USearch pipeline to process reads into OTU calls; the USearch sintax algorithm was used to designate the OTUs using the Ribosomal Database Project (RDP) – the reference database for 16S rRNA gene sequences. This allows calculation of the types of bacteria (referred to by OTU code; see supplemental table 5.1 for raw data including read counts), the bacterial community structure, relative, and absolute abundances (reported in supplemental table 2). qPCR was used to calculate the absolute number of bacterial cells in each sample by targeting the 16S rRNA gene using a Femto Bacterial DNA Quantification Kit (Zymo Research International). DNA sequencing was also carried out on a negative control (no contamination was found), and a positive control containing *Haemophilus* bacteria (OTU 2).

### Analytical methods and utilisation of comparative decay data

To test the similarity/dissimilarity of community structure and diversity between all samples where sequence data was successfully collected (i.e. hypothesis 1), we utilised a Principal Coordinate Analysis (PCoA) using the software PAST Version 4.13 (Hammer *et al*. 2001), which allows analysis of bacterial community structure while removing abundance biases.

The analysis investigating the relationship between the change in the bacterial community structure through time and sequence of morphological character loss (i.e. hypothesis 2) was undertaken using the high-resolution morphological character presence/absence data for amphioxus collected by Murdock *et al*. (in prep). This bacterial communities within the decay environment on specific sampling days were directly compared with this morphological character data (see supplemental table 3). To investigate the potential correlation between character loss and bacterial community structure, we utilised a further Principal Coordinate Analysis. As morphological character data in decay experiments is binary (presence/absence), we had to transform our bacterial data to be comparable, and so we converted the OTU data into presence/absence binary data for each sample. Spearman’s Rank correlations were utilised to statistically compare the ‘amphioxus decay space’ and the ‘bacterial community structure space’ within the PCoA, to investigate any potential link between changes in bacterial community structure and the timing of loss of morphological amphioxus characters during decay.

### Limitations of experimental design

It is important to highlight difficulties with collecting data for these types of experiments, the low replicate numbers presented here, and to frankly caveat the interpretation of our findings. We note that, although we utilised appropriate DNA extraction kits for our experiment, we had a high rate of DNA extraction failures (see supplemental table 1.1; failures highlighted in yellow). We had a 94% failure rate of extracting viable bacterial DNA from amphioxus carcasses buried in kaolinite and mica sediments. There were also a high proportion of failures in bentonite (failure rate = 58%). Furthermore, the high level of failure means that on some sampling days there were complete extraction failures (e.g. day 8 and 42), while on others there were a low number of successful replicates. This limits our testing of the hypothesis regarding bacterial community succession and anatomical decay (i.e. hypothesis 2) to samples utilising bentonite and reduce the temporal resolution of our results. DNA extraction failures are not uncommon when attempting to extract bacterial DNA from decaying carcasses, and this has been previously attributed to low microbial biomass (e.g. Metcalf et al. 2013). It is difficult to positively identify the cause of these DNA extraction failures, however, it is possible that variables (such as refreezing) may have contributed. However, this work serves as a proof of concept and the experimental design has been refined in our ongoing investigations.

## Results

Across all samples examined in this study, 575 bacterial OTUs were identified: 509 to phylum level, 375 to order level, and 196 to genus level (supplemental table 1.3). The samples varied in the total number of OTUs (average: 153 OTUs, range: 11-379; supplemental table 2.1).

Data from the PCoA analysis investigating similarity/dissimilarity of community structure (Fig. 1. a.; supplemental table 4) reveal several important findings pertaining to the bacterial community structures of inoculated and non-inoculated decaying carcasses. Firstly, inoculum created from sediment and seawater taken from the Tamar River estuary, UK, has a very different bacterial community structure to the native bacterial community found within a decaying amphioxus carcass. In fact, the community structure of the Tamar River inoculum is clearly distinct from all samples in this study, even samples where carcasses were decayed within inoculated ASW.

**Figure 1.**
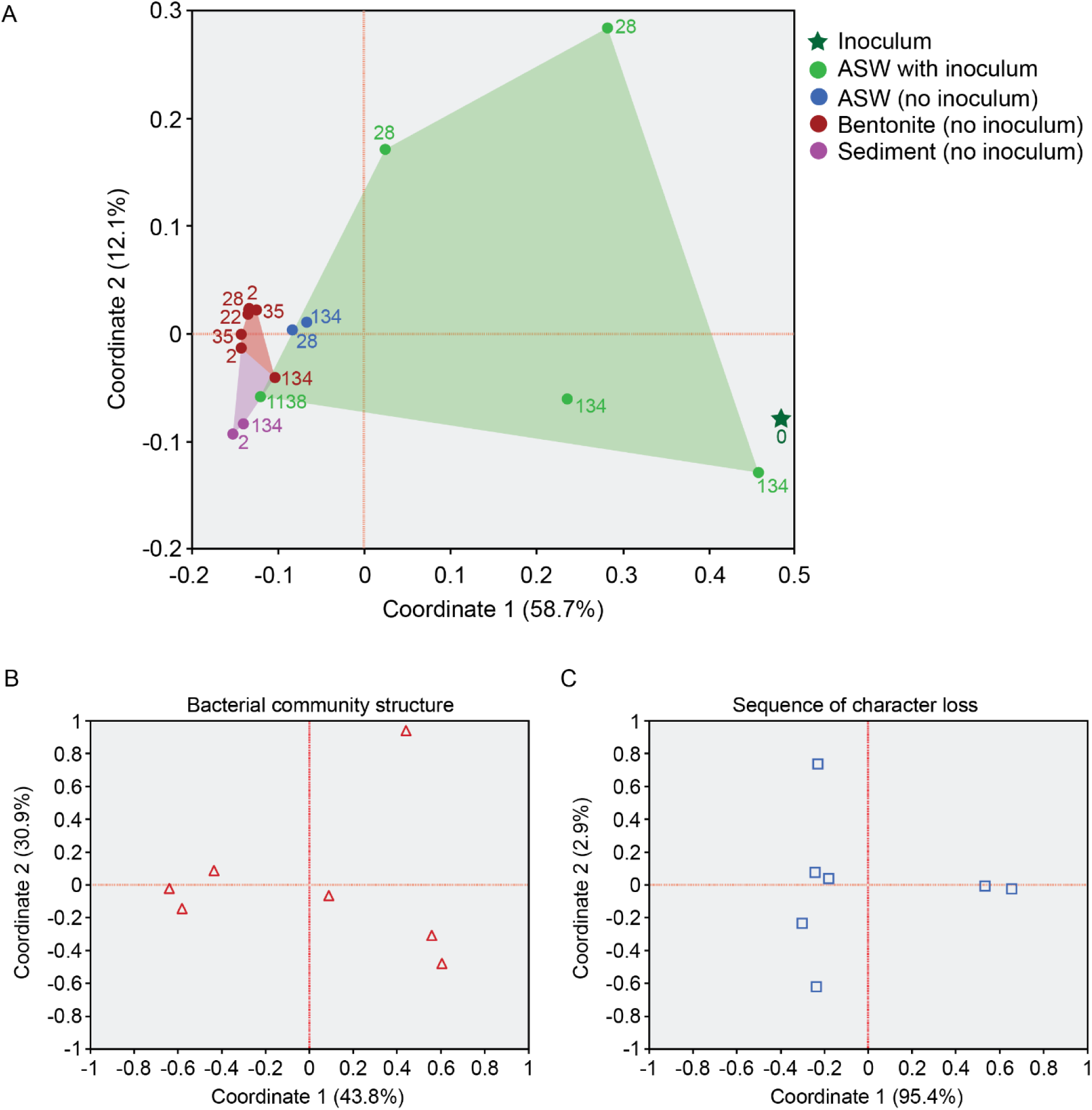
Analyses of bacterial communities within decaying amphioxi (*Branchiostoma lanceolatum*) carcasses. A, Principal Coordinate Analysis (PCoA) of presence/absence of bacterial OTUs in artificial seawater (ASW) controls, inoculated ASW samples, sediment samples, and pure ‘natural’ inoculum sample, to display the disparity of bacterial community structure between each sample. Purple samples represent non-bentonite sediment samples (kaolinite and mica). B, Principal Coordinate Analysis (PCoA) of presence/absence of bacterial OTUs in bentonite sediment samples, to display the disparity of bacterial community structure between each sample. C, PCoA of presence/absence of morphological characters within the decaying amphioxus carcass at each sampling point. Coordinates 1 and 2 are shown for each B and C. For eigenvalues, coordinate variance is represented as a percentage, and coordinate values, for all coordinates, see supplemental table 4.

Secondly, the data from the experiments designed to test hypothesis 1 show that decaying amphioxus carcasses within inoculated ASW had the largest data spread, indicating that the bacterial community varied in structure through time as decay progressed. The variability in bacterial community structure seen between the inoculated samples is greater than the variability between either samples decayed in ASW, or samples decayed within sediment. The data show the bacterial community structure within amphioxus carcasses that decayed in ASW (control sample) showed variability through time. In comparison, the bacterial communities within the amphioxus carcasses decaying in sediment show greater community structure variability through time than control samples, but far less than in those samples decayed in inoculated ASW. In terms of coordinate space, there is some minor overlap between the inoculated samples, and the non-inoculated samples - interestingly this overlap only occurs with the inoculum sample from day 1138. It should be noted that the non-bentonite sediment samples (amphioxus decayed in kaolinite or mica) do not overlap with the inoculated samples, however, this result should be interpreted cautiously as the high failure rate of DNA extraction in these sediments means there are limited replicates.

The experiment designed to test hypothesis 2 allowed investigation of the link between changes in bacterial community structure through time within a decaying amphioxus carcass entombed in bentonite (Fig. 1. b., Table 1, supplemental table 4) and the timing of the loss of Amphioxus’s anatomical characters during decay (Fig. 1. c., Table 1). Spearman’s Rank tests found a significant correlation between coordinate values on coordinate 2 (bacterial community structure) and time in Fig. 1. b., and between coordinate values on coordinate 1 (sequence of character loss) and organismal completeness in Fig. 1. c. (see also Table 1). However, our data show no correlation exists between coordinate values for coordinate 2 in Fig. 1. b. and coordinate 1 in Fig. 1. c. This result indicates that the change in bacterial community through time is not intrinsically linked to the sequence of anatomical character loss seen in a decaying amphioxus. This supports the null-hypothesis that character loss is controlled by the intrinsic anatomical structure of characters.

**Table 1:** Spearman’s rank correlations for PCoA coordinates of Fig. 1. b. in grey and Fig. 1. c. in white. The grey boxes represent tests comparing the presence/absence of bacterial OTUs in bentonite sediment samples against sample day (i.e. time). The white boxes represent tests comparing the presence/absence of morphological characters within the decaying amphioxus carcass at each sampling point with organismal completeness (see supplemental table 3). Results highlighted in red represent significant correlations. The yellow boxes represent tests comparing the coordinates displaying significant correlations between Fig. 1. b. and Fig.1. c. The data demonstrate no correlation between the change in the bacterial community structure through time (BCS) and the sequence of anatomical character loss (SCL).

| Experiment | Variable 1 | Variable 2 | Spearman's rho | p value |
| --- | --- | --- | --- | --- |
| Bacterial Community Structure | Coord 1 Bac Com | Time | -0.5637 | 0.1875 |
| Bacterial Community Structure | Coord 2 Bac Com | Time | 0.9456 | 0.0013 |
| Bacterial Community Structure | Coord 3 Bac Com | Time | -0.0909 | 0.8463 |
| Bacterial Community Structure | Coord 4 Bac Com | Time | -0.2546 | 0.5817 |
| Bacterial Community Structure | Coord 5 Bac Com | Time | -0.0182 | 0.9691 |
| Bacterial Community Structure | Coord 6 Bac Com | Time | -0.0364 | 0.9383 |
| Decay Sequence | Coord 1 Dec Seq | Completeness | 0.9636 | 0.0005 |
| Decay Sequence | Coord 2 Dec Seq | Completeness | 0.0741 | 0.8745 |
| Decay Sequence | Coord 3 Dec Seq | Completeness | 0.0371 | 0.9371 |
| Decay Sequence | Coord 4 Dec Seq | Completeness | 0.0385 | 0.9348 |
| BCS vs SCL Spearman's rank | Coord 1 Dec Seq | Coord 2 Bac Com | -0.6071 | 0.1482 |

### Bacterial succession dynamics within decaying amphioxus carcasses

The succession of bacterial communities through time within amphioxus carcasses decaying in inoculated and non-inoculated conditions are shown in Fig. 2. The data show that there is a succession of bacterial communities through time, although they are typically dominated by one or two OTUs.

**Figure 2.**
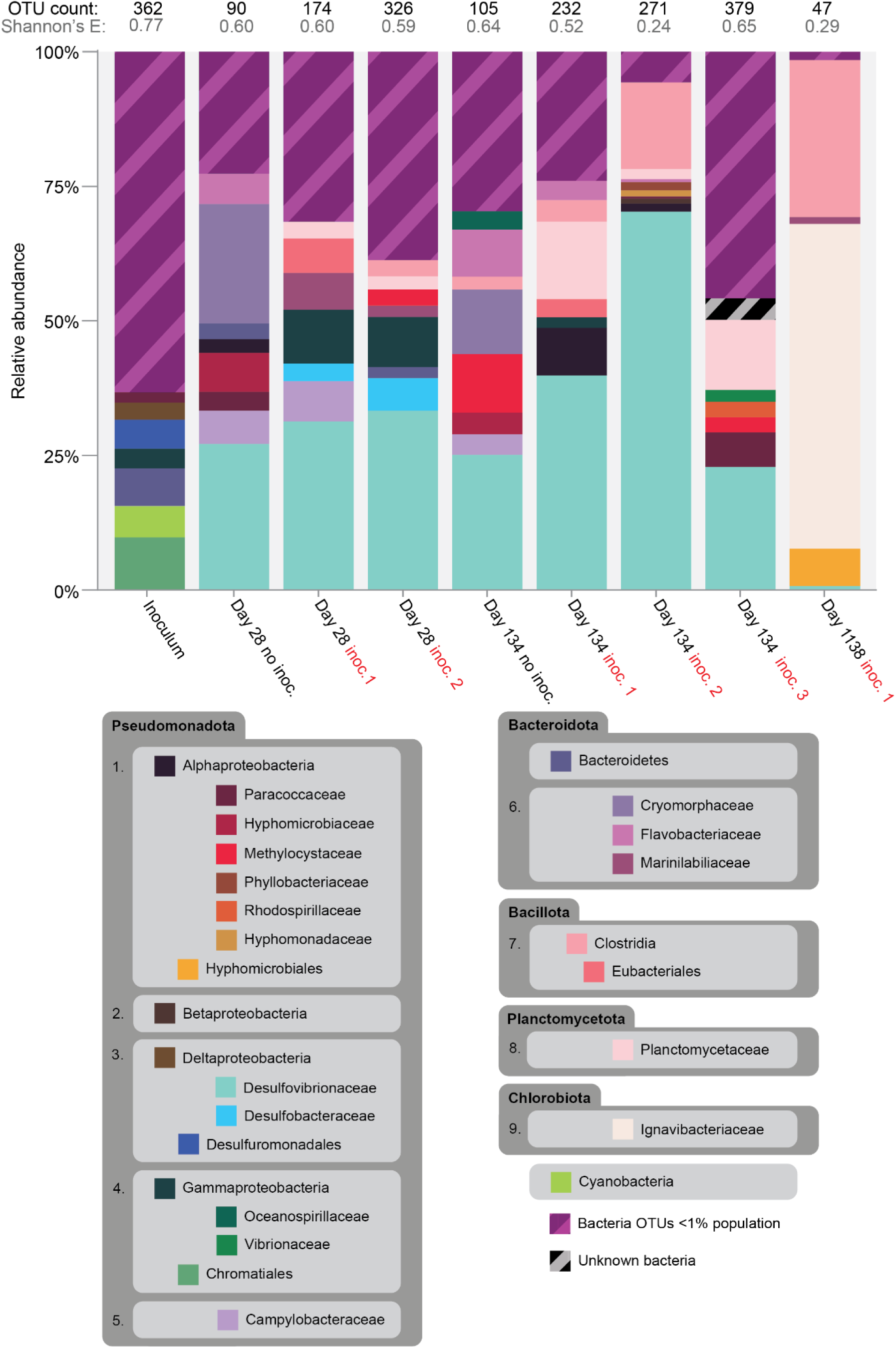
Relative abundance of dominant microbes within samples of pure inoculum and within decaying amphioxus (*Branchiostoma lanceolatum*) carcasses through time. Labelled bacteria in each sample represent bacteria where the relative proportion is greater than 1% of the population. All bacteria with relative abundances representing less than 1% of the community are agglomerated into the ‘Bacteria OTUs <1% population’ category. Bacteria taxonomy is reported to the family (or lowest possible) taxonomic rank based on Parte *et al*. (2020) and Oren and Garrity (2021). Sub-organised by class: 1) Alphaproteobacteria, 2) Betaproteobacteria, 3) Deltaproteobacteria, 4) Gammaproteobacteria, 5) Epsilonproteobacteria, 6) Flavobacteriia, 7) Clostridia, 8) Planctomycetia, 9) Ignavibacteria.

The pure inoculum has a diverse bacterial community with 363 identified OTUs and the highest Shannon’s equitability value of all samples (0.764, see Fig. 2; supplemental table 2.2). However, the majority of the OTUs identified represent tiny fractions of the overall community; 93% of OTUs each individually represent less than 1% of the relative abundance within the bacterial community. The inoculum’s bacterial community is dominated by Chromatiales, commonly known as purple sulphur bacteria (Hunter et al. 2009). Other major components of the bacterial community are Desulfuromonadaceae, Desulfuromonadales, Rhodobacterales, as well as OTUs representing Cyanobacteria, Bacteroidetes, Gammaproteobacteria, Bacteroidetes, Deltaproteobacteria although these OTUs cannot be identified beyond the order level.

The number of bacterial OTUs in the population of the day 28 decaying amphioxus carcass in a non-inoculated medium is smaller, with 90 OTUs identified - and only 17% of the OTUs individually represent a greater relative abundance than 1% of the bacterial community. The bacterial community structure differs from the inoculum, being dominated by Desulfovibrionaceae (OTU code 5; 27% relative abundance). At day 134 the number of OTUs in the bacterial community of the non-inoculated decaying amphioxus carcass is also smaller than the inoculum with 105 OTUs identified - and, again, only 17% of the OTUs have a greater individual relative abundance than 1% of the bacterial community. The bacterial community in this sample is also dominated by Desulfovibrionaceae (OTU code 5; 25% relative abundance). Both of the bacterial communities of these samples are secondarily dominated by Cryomorphaceae (22% and 11% respectively); however, the other dominant bacterial OTUs vary between samples, potentially indicating some disparate bacterial community structural changes through time.

Importantly, our data shows that, despite having much richer communities in terms of identified OTUs, the dominant OTU in all the inoculated samples is Desulfovibrionaceae (OTU code 5; day 28, 31% and 32%; day 134, 39%, 70%, and 21%). Desulfovibrionaceae (OTU code 5) was not identified in the inoculum at all, indicating that in samples containing decaying carcasses, this bacterial taxon represents a native OTUs from the carcass’s endogenous bacterial population. Comparatively, with the exception of the second replicate of day 134 (70% relative abundance), the relative proportion of Desulfovibrionaceae is similar to the non-inoculated samples (∼30%). Our data show that there are some bacterial community structure changes through time in the inoculated samples - samples on day 28 share high relative abundance of two Gammaproteobacteria taxa (OTU code 90 and 73), although neither of these bacteria have abundance above 1% in the day 134 samples. The bacterial communities compared between inoculated samples on day 134 are varied, sharing few dominant OTUs outside of Desulfovibrionaceae (OTU code 5).

OTU data was also collected on day 1138, well after the carcass had lost structural integrity and fully disarticulated. Despite having been inoculated at the onset of the experiment, the bacterial diversity on day 1138 was drastically different from all previous samples. The bacterial community was the smallest of any of the samples in this experiment stream with 47 identified OTUs and the lowest Shannon’s equitability value of the whole experiment (0.14; see Fig. 2, supplemental table 2.2). Proportionally, this sample is dominated by Ignavibacteriaceae (OTU code 434; 60%) and Clostridia (OTU code 28; 28%) - two bacterial taxa that do not present above 1% in any other previous sample. In fact, the day 1138 sample was dominated by four bacteria OTUs that had a relative abundance above 1%. The other 43 bacteria OTUs account for ∼3% of the relative abundance, and this includes Desulfovibrionaceae (OTU code 5) which only accounted for a relative abundance of 0.6%.

Calculating the structure of the bacterial communities also allows investigation into the major modes of bacterial respiration in each sample (Fig. 3). The bacterial community of the inoculum is dominated by bacteria that utilise anaerobic respiration (46%), but it also has the highest proportion of bacteria with an unknown mode of respiration (37%). The proportion of bacteria with an unknown respiratory mode is consistently higher in inoculated samples than in non-inoculated samples. Our data show clearly that all samples are dominated by bacteria that respire anaerobically (average: 71%). Further examination of the anaerobic bacteria data does indicate that, in the non-inoculated samples, greater proportions of the bacterial community’s diversity is made up of facultative anaerobes. This is not the case in the inoculated samples. Aerobically respiring bacteria are identified in all samples as small proportions of the community, although proportionally these communities are much lower in samples that were inoculated. The day 1138 sample is overwhelmingly dominated by anaerobically respiring bacteria (∼98%) with almost no aerobically respiring bacterial OTUs (> 1%) and a very low component of unknown bacteria (∼1%).

**Figure 3.**
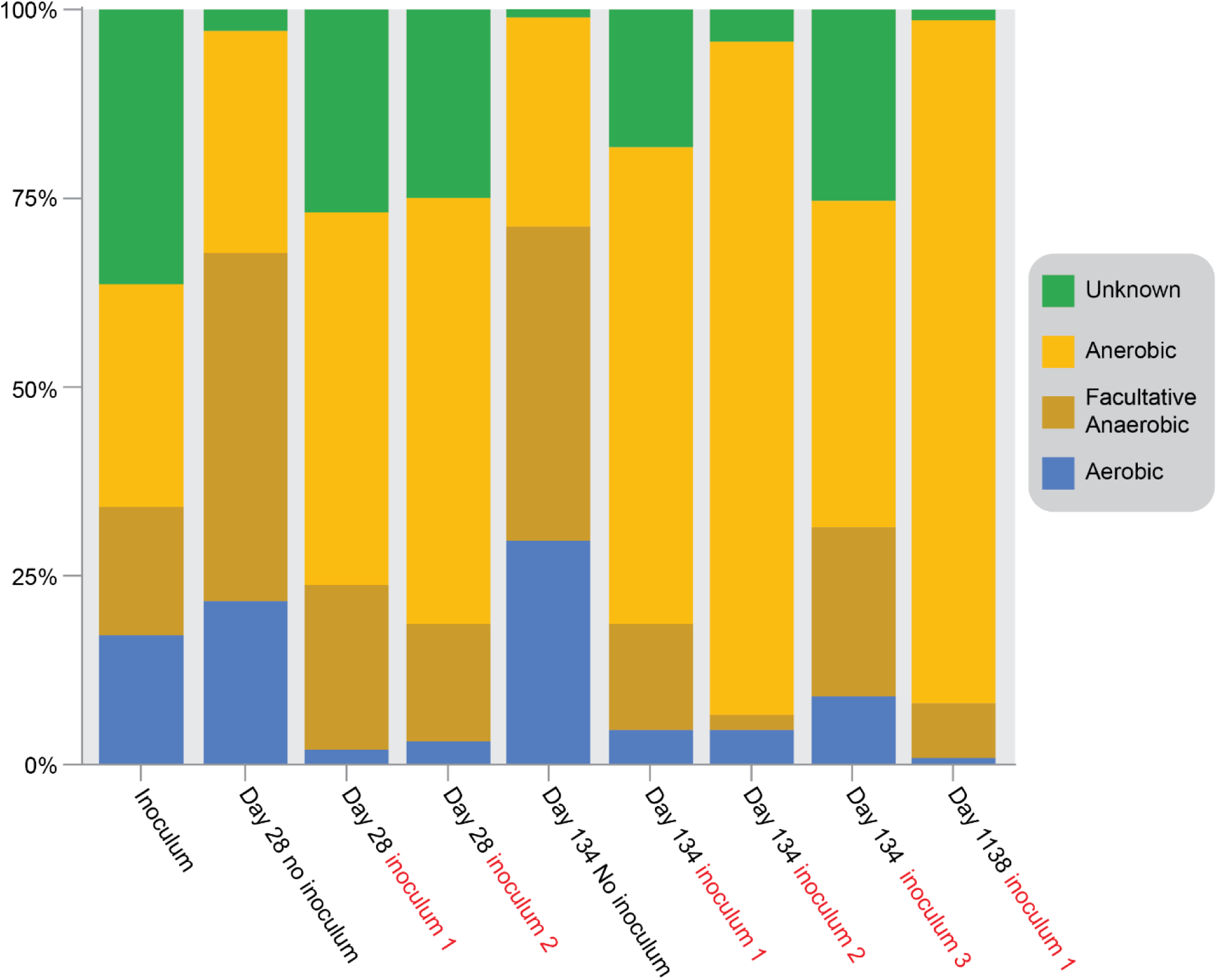
Relative proportions of the respiratory mode of the bacterial communities within decaying amphioxi carcass through time.

## Discussion

We are unable to reject our null hypothesis that the bacterial communities that develop within a decaying carcass through time are dominated by endogenous taxa; in both the inoculated samples and the control, the dominant bacterial OTUs within the thanatomicrobiome were not specific to, or found within the inoculum. The bacterial community of the inoculum was diverse, comprising in large part bacterial OTUs falling into the <1% abundance category, most of which are not found in either the control or inoculated samples. Adding an inoculum to decay experiments does not impact the endogenous thanatomicrobiome community structure, nor do exogenous bacteria seem to actively invade the carcass through the period of decay during which the carcass maintains its integrity. Our data show that in amphioxus carcasses, the thanatomicrobiome is typically dominated by the Desulfovibrionaceae: a family of anaerobic, sulphate-reducing bacteria, which are common components of the endogenous gut microbiota in a wide range of animals (e.g. Galushko and Keuever, 2020; Sayavedra et al. 2021; Singh et al. 2023). Our data, therefore, suggest that the endogenous anaerobic bacteria that inhabit an animal in life, proliferate and dominate the thanatomicrobiome within a decaying carcass (see next section for detailed description of thanatomicrobiome formation). This result has implications for the design of decay experiments. An exogenous inoculum added to a decay environment contains a large number of bacterial taxa with low abundance that are not persistent in the thanatomicrobiome during decay. Inocula are not required to initiate decay (e.g. Briggs and Kear 1993a, Sansom et al. 2010; Clements et al. 2017, 2021), so the addition of a natural inoculum to an experiment investigating anatomical decay contributes nothing more than an uncontrolled and potentially confounding variable. Without detailed microbial community analysis, the use of such inocula creates significant reproducibility issues; especially as bacteria are only a part of any natural inoculum taken from the wild: Fungi and Archaea may also be present. This, and our results, leads us to conclude that inocula are not recommended for most decay experiments.

We are also unable to reject our second null hypothesis, that the bacterial communities that develop within a decaying carcass through time are not correlated with the sequence of character loss. The community structure of the thanatomicrobiome is temporally dynamic through the decay process, but the lack of correlation indicates that, within a decaying carcass, it is not the succession of the bacterial communities that drives the patterns of anatomical character loss. But how far can we extend our results beyond amphioxus, and what do they mean for analysis of how non-biomineralised anatomy becomes fossilised? Gut microbiomes vary considerably between animals, even within a species (e.g. Luan et al. 2023; see refs in Davenport et al. 2017), posing the question of whether this might lead to variation in sequences of character loss? Current evidence suggests not because: (1) sequences of anatomical loss are conserved between individuals of the same species, and where homologues characters allow comparison, between species (Sansom et al 2010, 2011, Murdock et al 2014; Cavicchini et al. *in prep*); and (2) postmortem, there is significant microbial turnover (dysbiosis) in a carcass, with anaerobic bacteria, normally found in small numbers in the guts of animals (see below), coming to dominate the thanatomicrobiome within hours of death. Our current experiments are investigating this further, but considering our results here, microbial communities probably exhibit a high degree of convergence postmortem, particularly in terms of metabolic pathways to tissue degradation, suggested by the dominance of certain OTUs across almost all sampling intervals, despite the likely variation in initial endogenous microbiotas between specimens. Variation between living organisms’ endogenous microbiotas is thus likely to be significantly reduced after death.

These factors, combined with our results, have significant implications for the field of experimental decay and taphonomy: lack of knowledge about ancient microbial communities that caused decay and character loss in organisms in deep time does not matter. Bacterial successions are not correlated with anatomical character loss; non-biomineralised characters preserved in fossils correlate with their propensity to decay in extant relatives (e.g. Gabbott et al. 2021); the sequences of character loss are conserved between individuals and taxa (e.g. Sansom et al 2010, 2011, Murdock et al. 2014, Cavicchini et al. *in prep*) despite the likelihood of differences in gut microbiota between organisms in life. Together, this indicates that the specific taxa present in the microbiota is not likely to be a significant factor in controlling the loss of characters. Crucially, this supports the foundational hypothesis that anatomical decay is an experimentally tractable route to understanding how exceptionally preserved fossils form.

### Formation of the thanatomicrobiome, the microbial clock, and implication for soft tissue fossils

The concept of the thanatomicrobiome as a distinct internal microbiome within a decaying carcass is relatively new (Can et al. 2014). In forensic science, understanding the formation and succession of the thanatomicrobiome has attracted attention as a potential tool for criminal investigations, and more precise determination of ‘time of death’ or the postmortem interval (PMI) of cadavers than more traditional methods (see Javan and Finley, 2018; Metcalf, 2019; Deel et al. 2020). The rationale that postmortem interval can be resolved by investigation of the thanatomicrobiome comes from several studies that have identified a predictable ‘clock-like’ microbial community succession that occurs within a decaying carcass (e.g. Dickson et al 2011; Pechal et al. 2013; Metcalf et al 2013, 2016; Can et al., 2014; Cobaugh et al. 2015; Guo et al. 2016; Javan et al. 2016; Deel et al., 2020; Zhou et al 2021; Aragonés et al. 2022).

The potential importance of the thanatomicrobiome in a taphonomic context has not previously been considered explicitly. The link between bacteria and soft tissue preservation is well established, and we know that bacteria directly and indirectly influence preservational processes such as mineralisation (e.g. Sagemann et al. 1999; see review below), but what are the implications for the fossilisation of non-biomineralised anatomy? Unfortunately, the relatively high number of DNA extraction failures in our study limits the temporal resolution of our view of the formation and dynamics of the thanatomicrobiome immediately postmortem. Nevertheless, our data show that, even in the presence of an inoculum, the thanatomicrobiome is dominated by endogenous bacteria. How well does this accord with previous work on how the thanatomicrobiome develops and changes through time?

Before death, animals teem with bacteria; it is reported that the ratio of human cells to bacteria is approximately 1:1 (Sender et al. 2010; Walker and Hoyles 2023). In healthy living animals, while oral and skin microbiomes exist, only the guts and lungs have significant resident internal microbiotas. Most organs are considered to be largely sterile, with the immune system acting to restrict the organs and tissues that bacteria are able to inhabit (e.g. Can et al. 2014; Javan et al. 2016; Aragonés et al. 2022). Upon death, however, the immune system ceases to function and the mechanisms that control bacterial proliferation fail, allowing bacteria to rapidly colonise the carcass (Can et al. 2014; Aragonés et al. 2022). Where do these bacteria come from? In their review, Janssen et al. (2022) indicate that both endogenous and exogenous bacteria may contribute to decay and subsequent preservational processes, and while this is true, we note an important caveat. Exogenous bacteria may contribute to decay of the external surfaces of an animal carcass (these communities are known as epinecrotic bacterial communities) or create external biofilms (as per Eagen et al. 2017), but there is limited evidence to indicate that they invade and colonise the carcass, unless and until the carcass ruptures or is otherwise compromised. In fact, there is a growing body of evidence that, within an intact carcass, endogenous bacteria from the gut dominate the thanatomicrobiome postmortem. Several studies of mammalian carcasses (humans and mice) have documented this (e.g. Akutsu et al. 2012; Metcalf et al. 2013; Tuomisto et al. 2014; Javan et al. 2018), and Butler et al. (2015) observed that in *Artemia* (brine shrimp) gut bacteria proliferated and colonised the body cavity during decay. Our data provide additional support here. The dominant bacteria found in our samples, Desulfovibrionaceae, is a group of anaerobic bacteria found commonly in the guts of animals (e.g. Keuever 2014; Galushko and Keuever, 2015; Sayavedra et al. 2021).

During the first few hours of decay, the bacterial population within the gut is highly dynamic. High temporal resolution sampling of the thanatomicrobiome of decaying mice identified a distinct phase of gut bacterial community dysbiosis in the first 24 hours postmortem (Aragonés et al. 2022). In the first hours of decay, the ‘normal’ intestinal bacterial community collapses, probably due to the exhaustion of residual oxygen, and an ecological succession occurs where anaerobic bacteria outcompete and replace aerobic forms – this period, referred to as the adaptive phase, lasts for approximately 12 hours, until the anaerobic bacterial populations are established (Aragonés et al. 2022). Experiments on human cadavers have shown a similar pattern of postmortem gut bacteria dysbiosis, demonstrating that the thanatomicrobiome becomes dominated by a few genera (Metcalf et al. 2013; DeBruyn and Hauther 2017; Zhou et al. 2018). Our data supports this: the amphioxus thanatomicrobiome is rapidly dominated by the anaerobe group Desulfovibrionaceae. This finding is pertinent for soft tissue preservation because the rapid switch from aerobic to anaerobic as the main mode of bacterial respiration is a vital step in the generation of the geochemical conditions required for authigenic mineralisation to occur (e.g. Sagemann et al. 1999; Clements et al. 2017; 2022; see below).

For bacteria found in the gut during life to dominate the thanatomicrobiome, they must somehow proliferate throughout a carcass. During the decay of *Artemia*, Butler et al. (2015) reported that bacteria escaped from the gut when the organ began to lose structural integrity. Clements et al. (2022) similarly reported that during the decay of European sea bass (*Dicentrarchus labrax*) small ruptures occurred at the stomach and anal terminations which, theoretically, could allow gut bacteria to escape. While this is a plausible route for gut bacteria to colonise the body cavity, Clements et al. (2022) reported that the gut was one of the last organs to lose structural integrity with visible breakdown of the gut wall not occurring until the 11th day of decay. Therefore, it seems less likely that gut breakdown cannot account for the rapid and sequential manner of organ colonisation by anaerobic gut bacteria commonly seen in human cadavers (e.g. Janven et al. 2018; Zhou et al. 2018). An alternative explanation has been suggested: bacteria are typically highly motile (Janssen et al. 2022), and prior to the breakdown of the gut wall, as the immune system ceases to function, gut bacteria may colonise other organs via the capillaries of the vascular and lymphatic system (Noriko 1995; Janven et al. 2018; Dash and Das, 2020). Logically, this must occur rapidly, because organ decay typically begins within hours of death (e.g. Gill-King et al. 1997; Clements et al. 2022), but further detailed investigation is needed to establish empirically how anaerobic gut bacteria proliferate and become established throughout the carcass.

As decay proceeds, decay-related gases build up within the carcass. Commonly, this is most prominent in the gut region, presumably because anaerobic bacterial communities are quickly established in the gut, and the gut itself acts to restrict the movement of gases. Eventually, if gas cannot escape the gut can rupture, often with enough force to breach the integrity of the carcass (e.g. Metcalf et al. 2013; Clements et al. 2021; Aragonés et al. 2022). When this happens, exogenous bacteria can rapidly colonise and replace endogenous populations (e.g. Metcalf et al. 2013; Javan et al. 2018; Aragonés et al. 2022). The rapidity of this process makes mapping the bacterial community succession extremely difficult. Moreover, because the environmental conditions outside the carcass can vary across a broad range, exogenous bacterial colonisation is hard to predict, limiting its use as a tool for postmortem interval estimations (Lemon et al. 2012; Javan et al. 2018). Importantly, although many forensic studies of the ‘microbial-clock’ focus on this time interval during which rupture occurs, their experimental designs involve decaying human cadavers (or other mammals, often swine) in poorly constrained, terrestrial, subaerial environments. This lack of constraint makes robust and repeatable analysis of post-rupture bacterial successions difficult (see Aragonés et al. 2022). However, the many forensic studies showing that bacterial communities within ruptured carcasses rapidly become dominated by aerobic bacteria (e.g. Metcalf et al. 2013; Aragonés et al. 2022) are likely to have limited applicability to the kind of environments where exceptional preservation of non-biomineralised tissues takes place. In these environments, typically marine, with carcasses enclosed in sediments with limited pore water movement or oxygen exchange mechanisms, experiments show that decay-induced anaerobic conditions form rapidly (Sagemann et al. 1999) and that external geochemical gradients can often persist long after carcass rupture (Clements et al. 2022). In our experiment, we sampled on day 1138, long after the carcass lost its integrity (Fig. 2). In this sample, the thanatomicrobiome present in the previous samples had been replaced. The 1138 sample was almost completely dominated by anaerobic bacteria (Fig. 3), and of the two OTUs that dominate the sample, Clostridia are seen in the thanatomicrobiome of many earlier samples, but Ignavibacteriaceae - the most abundant in sample 1138, are not (Fig. 2). Unfortunately, our experimental design and the vast time gap between the samples (1004 days) mean we cannot say if the rupture of a carcass in a sediment-enclosed marine setting would lead to ingress of exogenous anaerobic bacteria and a dramatic community turnover, but it is possible. Further experiments are required to test this and understand the nature and succession of the bacterial communities that develop in close proximity to a decaying, sediment-enclosed carcass compared to those within.

### Review of the complex role of bacterial metabolism in soft tissue preservation

If an animal carcass is buried immediately postmortem in an environment amenable for fossilisation, what factors govern the likelihood of soft tissue preservation, and what role do bacteria play in these processes? Three principal factors govern soft tissue preservation: rate of decay, the sequence of anatomical character loss, and processes of preservation (we focus on mineralisation as it is the more common mode of preservation). All three of these factors are intrinsically linked to bacterial metabolism.

Almost immediately postmortem, the carcass begins to decay through a combination of autolysis and bacterial metabolism. Autolysis starts within minutes of death: the functional processes of cells cease, and endogenous enzymes begin to act on the components of each cell itself, effectively ‘self-digesting’ the cell and surrounding tissues (e.g. Hyun et al. 2012; Zapico et al. 2014; Guo et al. 2016). Autolysis alone would eventually break down a carcass, but it is relatively slow. Importantly, however, this process triggers blooms in bacterial communities that metabolise the amino acids, carbohydrates, lipids, water etc. that are released from the rupturing cells (e.g. Can et al. 2014). As bacterial communities become established throughout the carcass (see previous section), bacterial metabolism rapidly becomes the dominant driver of organic matter breakdown (Raff et al. 2008; Jassen et al. 2021). In concert, these two processes will break down the carcass, driving loss of anatomical information, disarticulation, and eventually, the complete destruction of organic matter. However, decay of organic material within a carcass is not random – experimental work has shown that anatomical characters decay in a repeatable sequence (Sansom et al. 2010). This is important, because the complex interplay between rate of decay and the sequence of anatomical character loss determines the organic template upon which preservational processes (e.g. mineralisation) can act (e.g. Purnell et al. 2018; Gabbott et al. 2021). It is also important to note here that while some organic tissues may be geologically stabilised, all other tissues that are not stabilised by a preservational process will continue to decay until obliteration, including biomineralised tissues (Clements and Gabbott, 2021; Gabbott et al. 2021; see Purnell et al. 2018 for review).

Rate of decay is influenced by environmental variables that predominantly moderate bacterial metabolism. Rapid sedimentation and conditions at the water-sediment interface commonly characterised as ‘inhospitable’ suppress the activity of scavengers and sedimentary bioturbators, protecting the carcass from rapid disarticulation during the early phases of decay. It is a common misconception that environmental factors such as low temperature, high salinity, low oxygen content etc. stop decay (Clements and Gabbott, 2021). In fact, these factors only slow the rate of decay through modulation of bacterial activity (Briggs and Kear 1993a; Briggs 2003; Muscente et al. 2023). Even anoxia does not halt decay: many bacterial groups have evolved diverse metabolic adaptations that allow them to exist in low oxygen environments. These bacteria utilise electron acceptors other than oxygen to allow respiration to continue through a variety of anaerobic metabolic pathways (Allison 1988b; Briggs and Kear, 1993a, b; Sagemann et al. 1999; Briggs 2003; Janssen et al. 2022). Even if burial occurs in an oxic sedimentary environment, with limited pore water movement (‘closed’ conditions) experiments show that the local supply of oxygen is rapidly depleted by bacterial respiration (e.g. Sagemann et al. 1999), forcing a swift transition in the dominant bacterial metabolic pathway from aerobic to anaerobic respiration (Briggs et al. 1993a, b; Sagemann et al. 1999; our results presented here).

Unless preservational processes act to convert or replace organic tissues into remains that are stable over geological timescales, decay will continue until complete carcass obliteration. What initiates preservation? A growing body of evidence suggests that, somewhat paradoxically, the activity of anaerobic bacterial metabolism may trigger and sustain the replacement/replication of soft tissues by authigenic minerals (minerals that precipitate and grow *in situ*; see Briggs, 2003). Anaerobic bacteria can induce mineralisation indirectly and/or directly (referred to as bacteria-induced mineral precipitation; e.g. Hoffmann et al. 2021). Indirect mechanisms include: 1) the breakdown of organics by bacteria to liberate ions from tissues, not only fuelling mineralisation through supersaturating the local environment, but also creating potential sites for mineral nucleation on decaying organic structures (e.g. Briggs and Kear 1993, 1994; Wilby and Briggs 1997; Clements et al. 2022; Janssen et al. 2022); 2) bacterial biofilms that form during decay acting to pseudomorph soft tissues and mediate mineralisation (e.g. Raff et al. 2013; Butler et al. 2015; Eagan et al. 2017) and; 3) anaerobic bacterial respiration and metabolism fundamentally altering the geochemistry within and around the carcass (e.g. Sageman et al. 1999; Clements et al. 2017; 2022). It is well understood that many modes of mineralisation, such as precipitation of calcium phosphate, pyrite etc., require specific geochemical conditions to trigger and/or maintain tissue replacement (e.g. Briggs and Wilby, 1996; Sagemann et al. 1999; Clements et al. 2017; 2022), and these are also influenced strongly by bacteria. Experimental work has revealed significant geochemical gradients within a decaying carcass (Clements et al. 2022), and to a lesser extent in the local sediment surrounding the carcass (Sageman et al. 1999; Clements et al. 2017; 2022), and the dramatic shift towards conditions amenable for mineralisation is linked to increasing amounts of waste products of bacterial respiration within the localised area, including CO_2_ (aq), sulfuric acid (H_2_SO_4_), and fatty acids (Sageman et al. 1999; McNamara et al. 2009; Clements et al. 2017; 2022). In solution, these chemicals create intense localised geochemical gradients around decaying organics which can actively promote authigenic mineral precipitation (e.g. Sagemann et al. 1999; Clements et al. 2017; 2022). Interestingly, the generation of reducing conditions has been shown to also suppress the impact of autolysis, retarding the rate of decay (Raff et al. 2006; Butler et al. 2015). This may prolong the window for mineralisation to occur. Burial also plays an important role here: if the carcass is entombed, a closed environment is created with the sediment acting to limit diffusion of decay products away from the carcass, thus prolonging the duration of the intense local geochemical gradients caused by decay and increasing the likelihood that mineralisation can occur (Sagemann et al. 1999; McCoy et al. 2015; Clements et al. 2017).

Mineralisation is also restricted both temporally and spatially during decay; even if geochemical controls are amenable, modes of mineralisation are discriminate towards specific tissues within a carcass (e.g. Gabbott et al. 2021; Clements et al. 2022). This can be seen in the fossil record: patterns of soft tissue preservation clearly show that modes of mineralisation are biased towards the preservation of specific tissues and organs (e.g. Wilby, Briggs and Riou, 1996; McNamara et al. 2009; Jauvion et al. 2020; for a review see Clements et al. 2022). This has been explained by hypotheses of organ specific microenvironments (e.g. McNamara et al. 2009), but experimental investigations to date have failed to find evidence for such microenvironments. Rather, pervasive geochemical environments develop throughout a carcass, suggesting that once the geochemical conditions required for mineralisation are established, it is the intrinsic biochemical and structural properties of tissues that governs the likelihood of mineral replacement (see Clements et al. 2022).

Bacterially induced mineralisation reflects a delicate balance between competing processes. While bacterial metabolism is required to generate the geochemical conditions that allow mineralisation to occur, too much metabolism will obliterate soft tissues, removing the anatomical information that defines exceptional preservation of non-biominealised remains (Clements and Gabbott, 2021). Clearly, as soft tissue remains in the fossil record are so rare, minute variations in any of these variables may prevent soft tissue preservation from occurring. Decay experiments have improved our understanding of how non-biomineralised tissues become fossilised dramatically in the last three decades, but many aspects still remain poorly understood – and while fossil material gives us direct evidence of the end results, it is detailed and robust interdisciplinary experimental work that will provide the insights we need to address these gaps in our knowledge.

## Conclusion

Bacteria play a direct and indirect role in influencing preservational processes that geologically stabilise non-biomineralised tissues. Despite this importance, it has not been clear endogenous or exogenous bacteria dominate the bacterial communities that inhabit a carcass immediately postmortem (the thanatomicrobiome). Furthermore, recent work in microbial forensic science has suggested that successional shifts in the thanatomicrobiome may control the reproducible sequence of anatomical character loss seen in decaying carcasses. Here, we investigate the succession of the thanatomicrobiome within decaying amphioxus carcasses and present two key findings: firstly, even in the presence of inoculum, the thanatomicrobiome is dominated by endogenous anaerobic bacteria sourced from the digestive tract, and that exogenous bacteria do not invade the carcass and replace native bacteria species while the carcass is intact. Our data confirm previous findings that adding an inoculum to a taphonomic experiment is not required for decay to occur, and without detailed microbial, Archean, and fungal community analysis, naturally sourced inocula represent an uncontrolled and potentially confounding variable. Secondly, our data does not support that temporal successions in the thanatomicrobiome control the sequence of anatomical character loss during decay. This further supports the hypothesis that non- biomineralised characters preserved in fossils correlate with their propensity to decay in extant relatives. Furthermore, when combined with the evidence that the sequence of anatomical character loss during decay is conserved between individuals and across non-related taxa, we conclude that the inability to model ancient bacteria communities does not invalidate decay experiments. Our results provide a valuable ‘proof-of-concept’ which we are building upon, and our further experimental work will address the gaps in our understanding of the formation and proliferation of the thanatomicrobiome during the early stages of decay.

## Acknowledgments

Hector Escriva (Observatoire Océanologique de Banyuls-sur-Mer) is thanked for providing amphioxus and hospitality in Banyuls sur Mer. Thanks to Dr Sylvius from the Genomics Core Facility of the University of Leicester for support with the Illumina sequencing. A. Clements and E. Dunne are thanked for proof reading and feedback. Thomas Clements was funded by a Leverhulme Early Career Fellowship (ECF-2019-097). Purnell, Gabbott, Murdock, and Goodall were funded by NERC Grant NE/K004557/1 to MAP and SG. For the purpose of open access, the author has applied a Creative Commons Attribution (CC BY) licence to the Author Accepted Manuscript version arising from this submission.

## Contributions Statement

**Conceptualization** Mark Purnell (MAP), Sarah Gabbott (SG), Duncan Murdock (DJM); **Data Curation** Robert Goodall (RHG), Christopher Turkington (CT); **Formal Analysis** RHG, Thomas Clements (TC), MAP; **Funding Acquisition** MAP, SG; **Investigation** RHG, MAP, DJM, SG, CT; **Methodology** MAP, SG, DJM, RHG, Martha Clokie (MC), Andrew Millard (AM); **Project Administration** DJM, RHG, MAP; **Resources** Hector Escriva (Observatoire Océanologique de Banyuls-sur-Mer); **Supervision** MAP, SG; **Validation** TC; **Visualization** TC, RHG, MAP; **Writing – Original Draft Preparation** TC, RHG, MAP; **Writing – Review and Editing** TC, MAP, Orla Bath Enright (OBE), RHG, SG, DJM, AM, CT.

## Data availability

Until publication, supplementary data files are available here: https://doi.org/10.17605/OSF.IO/9CJ8M

Upon publication, supplementary data will be available on journals preferred repository.

## Notes

https://doi.org/10.17605/OSF.IO/9CJ8M

### Competing Interest Statement

The authors have declared no competing interest.

